# Serotonin therapies for opioid-induced dysphagia and respiratory depression: sex differences in a rat electromyography model

**DOI:** 10.1101/2023.08.21.554164

**Authors:** Michael Frazure, In Morimoto, Nathan Fielder, Nicholas Mellen, Kimberly Iceman, Teresa Pitts

## Abstract

Opioids are well-known to cause respiratory depression, but despite clinical evidence of dysphagia, the effects of opioids on swallow excitability and motor pattern are unknown. We sought to test the effects of the clinically-relevant opioid buprenorphine on pharyngeal swallow and respiratory drive in male and female rats. We also evaluated utility of serotonin 5-HT_1A_ agonists (8-OH-DPAT and buspirone) to improve swallowing and breathing outcomes following buprenorphine administration. Experiments were performed on 44 freely breathing Sprague Dawley rats anesthetized with sodium pentobarbital. Bipolar fine wire electrodes were inserted into the mylohyoid, thyroarytenoid, posterior cricoarytenoid, thyropharyngeus and diaphragm muscles to measure electromyographic (EMG) activity of swallowing and breathing behaviors. We evaluated the hypotheses that swallow varies by stimulus, opioids depress swallow and breathing, and that 5-HT_1A_ agonists improve these depressions. Our results largely confirmed the hypotheses: 1) Swallow-related muscle activity was larger during swallows elicited by oral water infusion plus esophageal distension than by either stimulus alone. 2) Buprenorphine depressed swallow in both sexes, but most significantly in females. 3) Female animals were more susceptible to buprenorphine-induced respiratory arrest. 4) 8-OH-DPAT rescued breathing following buprenorphine-induced respiratory arrest, and pre-treatment with the partial 5-HT_1A_ agonist buspirone prevented buprenorphine-induced respiratory arrest in female animals. 5) 8- OH-DPAT enhanced swallow-related mylohyoid drive, but did not restore excitability of the swallow pattern generator following total suppression by buprenorphine. Our results highlight sex-specific and behavior-specific effects of buprenorphine and provide pre-clinical evidence of a 5HT_1A_ agonist for the treatment of respiratory depression and dysphagia.

**New & Noteworthy:** This is the first manuscript to evaluate sex-specific effects of opioid administration on pharyngeal swallow. We expand on a small but growing number of studies that report a lower threshold for opioid-induced respiratory depression in females than males, and are the first to produce this effect with the partial mu-opioid-receptor agonist buprenorphine. Our study is the first to demonstrate that activation of 5-HT_1A_ receptors can improve swallow and breathing outcomes following systemic buprenorphine administration.

## Introduction

Precise coordination of the upper aerodigestive tract is essential for functional swallowing, breathing, and airway protection [1-3]. Swallow is the critical mechanism of energy intake for mammals [4], while breathing ventilates the lower airways to permit gas exchange [5]. Several protective reflexes, including swallow, prevent ingested food and liquid from compromising the airway [6-8]. Functionality of the upper aerodigestive tract may be disrupted by numerous disease states and drugs, including opioids [9-11]. The objective of our study was to systematically evaluate the effects of opioids on the upper aerodigestive tract and its dual role in breathing and swallowing.

Airway protection during swallow is achieved by synchronized laryngeal elevation and closure while ingested material is propelled through the pharynx [6]. Activation of pharyngeal mechanoreceptors triggers transient relaxation of the upper esophageal sphincter, enabling ingested material to pass into the esophagus [4]. Tonic at rest, the upper esophageal sphincter forms a functional barrier between the airway and any food contents in the esophagus [4, 7]. Failure to maintain separation of the airway from ingested material can result in aspiration of food or liquid into the lower airway [2, 3]. Sequelae of aspiration can be fatal if airway obstruction (acute) or aspiration pneumonia (chronic) occur [12].

The upper aerodigestive tract is active during both the inspiratory and expiratory phases of the respiratory cycle [2, 13, 14]. During inspiration, the glottis is abducted by the posterior cricoarytenoid muscle, and the upper esophageal sphincter is tonically contracted. In this configuration, inspired air meets less resistance from the larynx than from the upper esophageal sphincter, and flows through the glottis to the lower airways for gas exchange [4, 13]. During expiration, the larynx is partially adducted by the thyroarytenoid muscle. Partial laryngeal adduction functions as an expiratory braking mechanism that helps match ventilation to metabolic demand [2, 15]. The pharyngeal musculature can be active during either phase of the respiratory cycle, but is generally active during expiration [15].

Opioid receptors are inhibitory G-Protein Coupled Receptors (GPCRs), and are distributed throughout the pontomedullary respiratory network [16-22]. It is therefore unsurprising that respiratory depression is a serious complication of opioid use [23]. The United States is facing a distressing opioid epidemic: Opioid-related deaths have increased steadily for two decades, peaking each year since 2019 [23-25]. To complicate matters, adverse effects of opioid use are not limited to overdose events. Respiratory depression can also occur when therapeutic doses of opioids are administered in a controlled setting [23, 26-28].

In addition to well-known respiratory-depressant effects, opioids also depress the immune system, gastrointestinal system, and airway defense mechanisms (e.g., cough) [23, 29, 30]. Clinical trials have reported esophageal dysfunction and aspiration in humans following opioid administration, and several reports have identified aspiration pneumonia as a serious complication of opioid use following both overdose and chronic use [31-37]. While it is likely that aerodigestive dysregulation is a contributing factor to aspiration pneumonia following opioid use, specific effects of opioids on pharyngeal swallow have been the subject of limited investigation.

Buprenorphine is a partial μ-opioid receptor agonist that is used for post-operative pain management and opioid maintenance therapy [38-40]. Buprenorphine binds the μ-opioid receptor with high affinity for long durations, but as a partial agonist, activates the receptor to a lesser extent than a full agonist (e.g., morphine, fentanyl)[23, 39]. Because of these pharmacological properties, buprenorphine is generally considered to be safer than full μ-opioid agonists [41]. Indeed, a ceiling effect of buprenorphine-induced respiratory depression has been demonstrated in humans, but its effects on airway protection and deglutition are largely unknown [42, 43]. In the present study, we evaluated swallow and breathing function before and after buprenorphine administration to determine how a widely prescribed, clinically relevant opioid impacts aerodigestive function.

We initially tested three hypotheses: 1) Swallow motor pattern is modulated by aerodigestive afferent input to the medullary swallow pattern generator. 2) Administration of the opioid buprenorphine will result in a measurable decline of swallow function due to central depression of the swallow pattern generator. 3) Respiratory depression following systemic buprenorphine is not dose-dependent. During our initial dose response experiments, we found that, unlike males, most females succumbed to respiratory arrest following high doses of buprenorphine. This striking sex difference led us to consider potential counters to the potent respiratory depression we observed in female animals.

Additionally, basic science studies have implicated serotonin, a modulatory neurotransmitter, in the regulation of breathing and swallow [44-48]. Like opioid receptors, serotonin receptors are also GPCRs and are expressed throughout the brainstem swallow-breathing network [49]. Previous studies have demonstrated that the 5-HT_1A_ agonist 8-OH-DPAT produced excitatory respiratory effects in rats and rabbits [50, 51]. Notably, Sahibzada and colleagues reversed morphine-induced apnea in male rats using a 5-HT_1A_ agonist [52]. These promising reports, along with our primary results, led us to test three additional hypotheses: 4) Systemic administration of 8-OH-DPAT will restore breathing following buprenorphine-induced apnea through activation of 5-HT_1A_ receptors in female rats; 5) Pre-treatment with oral buspirone will preserve breathing following buprenorphine administration through action on 5-HT_1A_ receptors in female rats; 6) Systemic administration of a 5-HT_1A_ agonist will measurably improve swallow function following buprenorphine-induced depression of swallow

## Materials and Methods

### Study Design

All experiments were approved by the Institutional Animal Care and Use committee of the University of Louisville and conducted in accordance with the American Physiological Society’s Animal Care Guidelines [53]. Experiments were performed using 44 adult Sprague Dawley ex-breeder rats [21 male (0.58 ± 0.16 kg) and 23 female (0.26 ± 0.19 kg)]. Our objectives were to evaluate 1) differential effects of pharyngeal and esophageal stimulation on oropharyngeal swallow initiation and motor pattern, 2) how protective responses to upper aerodigestive stimuli (e.g., swallow) are affected by systemic opioid administration, and 3) the utility of 5-HT_1A_ agonists in the recovery/protection of swallow and respiratory drive following systemic opioid administration in rats.

### Rat Model

Animals were initially anesthetized with gaseous isoflurane (1.5% with 100% O_2_) while a femoral intravenous (IV) cannula (0.6 mm inner diameter) was placed. Animals were then transitioned to sodium pentobarbital (initial dose 25 mg/kg IV), with supplementary doses (1-4 mg/kg IV) administered as needed. A dose of atropine sulfate (0.01 mg/kg IV) was given at the beginning of each experiment to reduce airway secretions from repeated tracheal stimulation. Following administration of atropine sulfate, a tracheostomy was performed. Body temperature was monitored and maintained at 36.5 ± 0.5° C with a heating pad (Homeothermic Monitor, Harvard Apparatus). Anesthetic level was evaluated by jaw tone, blink reflex, forelimb withdrawal reflex, and licking in response to oral water administration.

### Electrophysiology Recording

Muscle activity was recorded via electromyography (EMG) using bipolar fine wire hook electrodes (A-M Systems stainless steel No. 791050) according to the technique of Basmajian and Stecko [55]. Four muscles were used to evaluate swallow: mylohyoid (hyolaryngeal elevator), thyroarytenoid (laryngeal adductor), thyropharyngeus (pharyngeal constrictor), and costal diaphragm. These muscles span the actions of the pharyngeal phase of swallow. Diaphragm activation during swallow (Schluckatmung) produces negative intrathoracic pressure and is innervated by the phrenic nerve (C3-5) [56-59]. The anatomical location of each muscle, and representative traces of muscle activity during pharyngeal swallow, are shown in Figure 1. Breathing motor patterns were evaluated using the posterior cricoarytenoid (laryngeal abductor, CN X), thyroarytenoid (laryngeal adductor), thyropharyngeus, and costal diaphragm [13, 15, 60]. Respiratory phase activity of inspiratory and expiratory upper airway muscles is shown in Figures 1 and 2.

**Figure 1.**
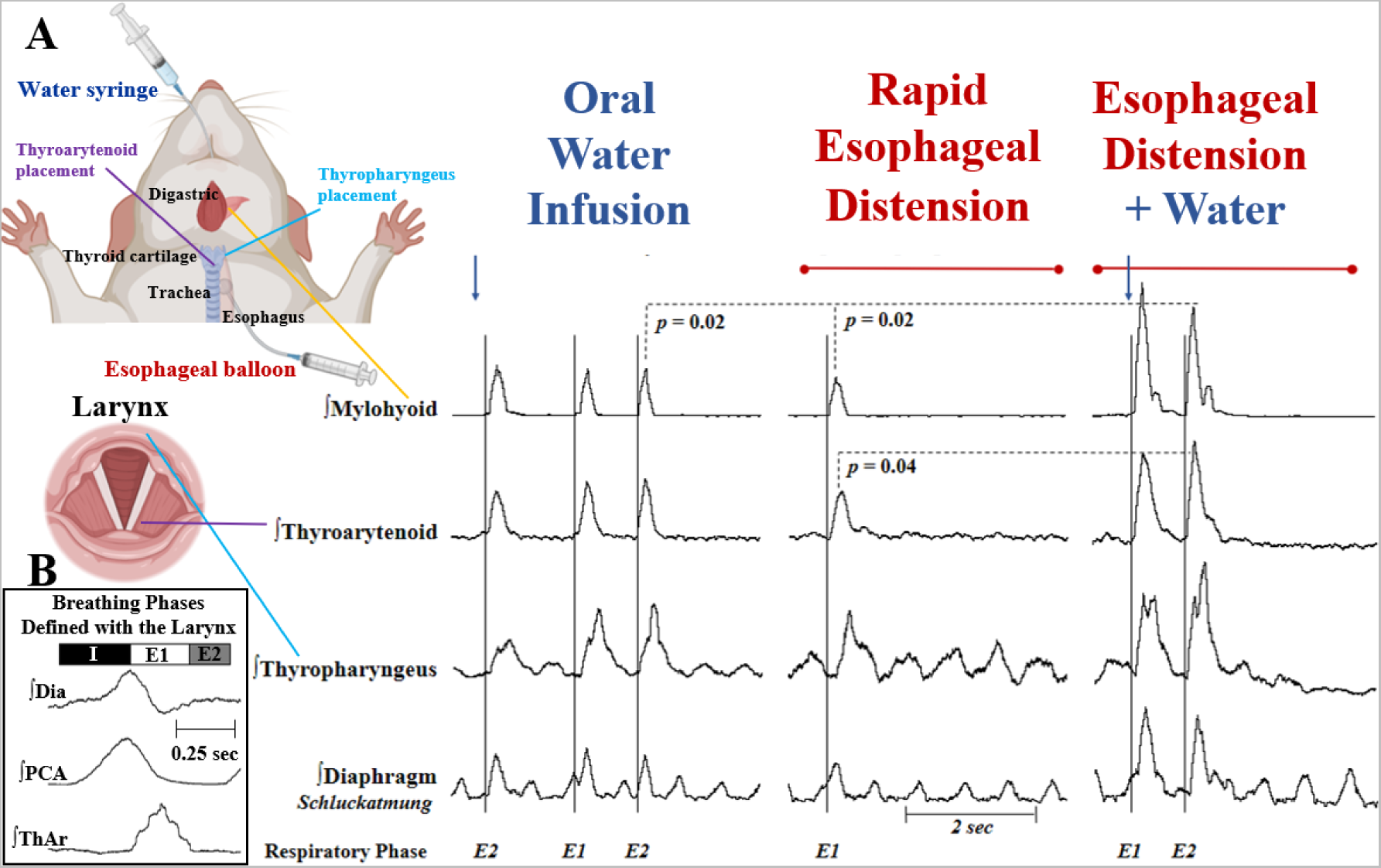
Simultaneous activation of pharyngeal and esophageal mechanoreceptors increases laryngeal drive during swallow. A) Electromyogram (EMG) activity was recorded from the mylohyoid, thyroarytenoid, thyropharyngeus and costal diaphragm muscles in freely breathing (pentobarbital anesthetized) rats with intact vagi. Swallows were elicited with infusion of 1 cc of water into the oropharynx, rapid upper esophageal distension (0.5 cc balloon volume), and combined stimulus (esophageal distension plus water infusion). Traces are rectified and integrated (20-ms), and amplitudes are reported as percent of mean. Arrows indicate water infusion in the oropharynx, horizontal lines indicate esophageal distension, and vertical lines indicate swallow initiation. Representative traces of EMG activity during swallow show stimulus-dependent modulation of the swallow motor pattern. Analysis of variance (ANOVA) showed differences in mylohyoid (laryngeal elevator) amplitude [*F*_(1.8, 40)_ = 7.93, *p* = 0.002], mylohyoid burst duration [*F*_(2.0, 27)_ = 14.19, *p* < 0.0001], and thyroarytenoid (laryngeal adductor) amplitude [*F*_(1.4, 31)_ = 4.19, *p* = 0.037] during swallow across the three stimulus conditions. Post-hoc comparisons using the Tukey HSD test indicated that the mean mylohyoid amplitude was larger during swallows elicited by esophageal distension plus water (118% ± 17) compared to swallows elicited by water alone (94% ± 19) or esophageal distension alone (92% ± 25). Mean mylohyoid burst duration was longer during swallows elicited by esophageal distension (304 ms ± 70, *p* = 0.0003) and esophageal distension plus water (311 ms ± 69, *p* = 0.0009) compared to swallows elicited by water alone (243 ms ± 58). Mean thyroarytenoid amplitude was larger during swallows elicited by esophageal distension plus water (104% ± 13) compared to swallows elicited by esophageal distension alone (91% ± 10). Diaphragm EMG activity reflects inspiratory activity during breathing and swallow. In 50% of animals, amplitude of diaphragm activity during swallows (i.e., Schluckatmung) elicited by esophageal distension and esophageal distension plus water increased qualitatively compared to swallows elicited by water alone, but the effect was not significant as a group. Respiratory phase was determined by diaphragm activity at the onset of swallow. Most swallows occurred during expiration, with no significant change in swallow-breathing coordination across stimulus conditions. B) Representative EMG example of breathing using laryngeal drive to define breathing phases. Traces are rectified and integrated (80 ms time constant). The diaphragm (Dia) acts as an inspiratory pump. The inspiratory-phasic posterior cricoarytenoid (PCA) opens the glottis, which reduces resistance to inspired air. Activation of the thyroarytenoid (ThAr) during expiration partially closes the larynx, and functions as a braking mechanism for the expiratory phase. Inspiration (I) is the period from the onset of diaphragm and PCA activation to the peak of the diaphragm burst. Early expiration (E1) is the period from the peak of the diaphragm burst to quiescence of the thyroarytenoid. Late expiration (E2) is the period from the end of the thyroarytenoid activation to the beginning of the next diaphragm burst. Breathing phases are defined by the larynx during eupnea. Because the thyroarytenoid is active during swallow, swallow-breathing coordination may be defined by diaphragm EMG activity.

**Figure 2.**
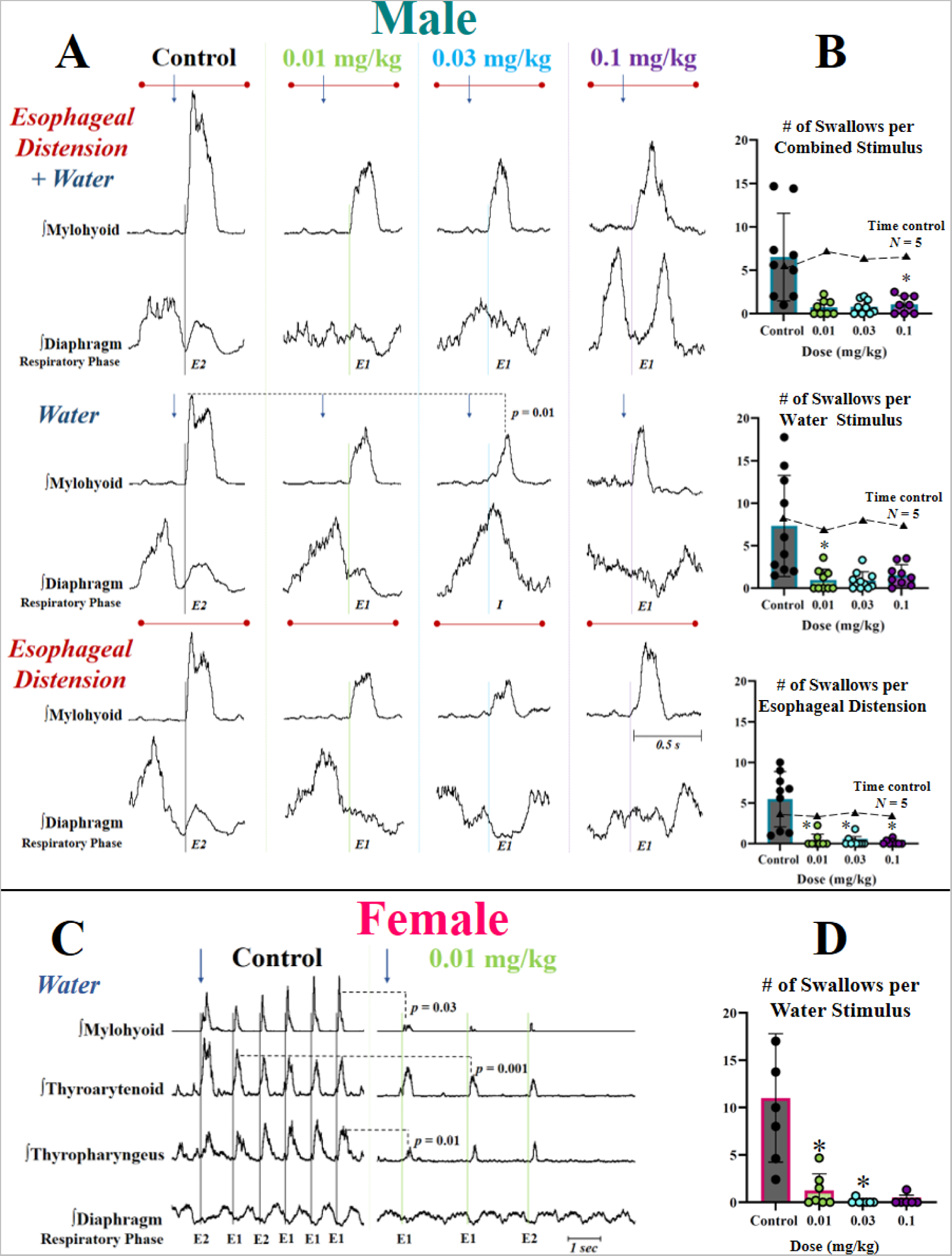
The opioid buprenorphine depresses swallow-related trigeminal drive in male rats, and swallow-related trigeminal, vagal and glossopharyngeal drive in female rats. To test the hypothesis that opioids depress swallow function, we performed cumulative dose response experiments with buprenorphine (0.01, 0.03 and 0.1 mg/kg IV) in freely breathing (vagi intact) pentobarbital anesthetized male and female Sprague Dawley Rats. A) Buprenorphine reduces laryngeal elevation during swallow, and alters swallow breathing coordination in male rats. Representative electromyogram (EMG) traces show mylohyoid (laryngeal elevator, innervation: CN V) and diaphragm muscle activity during swallows elicited by esophageal distension plus water (top), water alone (middle) and esophageal distension alone (bottom) across buprenorphine doses in a male rat. Traces are rectified and integrated (20 ms time constant), and amplitudes are reported as percent of mean during control. Arrows indicate water infusion in the oropharynx, horizontal lines indicate esophageal distension, and vertical lines indicate swallow initiation. All three stimuli reliably elicit swallow, and analysis of variance (ANOVA) showed significant differences in mylohyoid amplitude [*F*_(2.2, 9.7)_ = 7.3, *p* = 0.01] and burst duration [*F*_(1.3, 6)_ = 6.194, *p* = 0.04] during swallow. Post-hoc comparisons using the Tukey HSD test indicated that peak mylohyoid amplitude was significantly reduced during swallows elicited by water after 0.03 mg/kg buprenorphine (43% ± 15) compared to control water swallows (93% ± 19). A trending reduction in peak amplitude during swallows elicited by esophageal distension plus water after 0.01 mg/kg (57% ± 13, *p* = 0.09) and 0.03 mg/kg buprenorphine (72% ± 33, *p* = 0.06) compared to control combined stimulus swallows (115% ± 20) did not reach significance. Mylohyoid burst duration was significantly reduced during swallows elicited by water after 0.01 mg/kg (*M* = 196 ms ± 84) and swallows elicited by esophageal distension plus water after 0.03 mg/kg buprenorphine (170 ms ± 103), compared to control water (269 ms ± 56) and combined stimulus (*M* = 337 ms ± 44) swallows, respectively. Swallow breathing coordination was determined by respiratory phase at time of swallow onset. Swallow initiation was categorized as occurring during inspiration (I), early expiration (E1), or late expiration (E2). During control, most swallows occurred during late expiration. A Wilcoxon signed-rank test detected a significant change in swallow breathing coordination following buprenorphine administration, with significantly more swallows occurring during early expiration following buprenorphine administration, across dose and stimulus conditions (*z* = -3.71, *p* < 0.0001). B) Buprenorphine reduces frequency of swallow occurrence in male rats. Plots show number of swallows elicited by water plus esophageal distension, water, and esophageal distension stimuli before and after buprenorphine administration in male rats. Each circle represents the average number of swallows per stimulus for an individual animal across the buprenorphine dose response curve. Triangles represent the pooled average number of swallows elicited by combined stimulus, water, and esophageal distension during separate time control experiments (*n* = 5) in which animals received a saline vehicle infusion (0.5 cc IV) instead of buprenorphine at each time point of the dose response. ANOVA showed a significant difference in frequency of swallow occurrence [*F*_(1.8, 13.6)_ = 11, *p* = 0.002] among the buprenorphine experiments. Post-hoc comparisons using the Tukey HSD test showed that significantly fewer swallows occurred following: Oral water infusion and esophageal distension following 0.01 mg/kg buprenorphine; esophageal distension following 0.03 mg/kg buprenorphine; and esophageal distension plus water and esophageal distension alone after 0.1 mg/kg buprenorphine, compared to control, respectively. ANOVA showed no significant change in frequency of swallow occurrence among time control animals [*F*_(1.8, 7.3)_ = 3.3, *p* = 0.1]. * indicates *p* < 0.05. C) Buprenorphine reduces laryngeal elevation, laryngeal closure, and pharyngeal constriction during swallow in female rats. Representative EMG traces of mylohyoid (laryngeal elevator, innervation: CN V), thyroarytenoid (laryngeal adductor, innervation: CN X), thyropharyngeus (pharyngeal constrictor, innervation: CN IX) and diaphragm depict swallows elicited by oral water infusion before and after buprenorphine administration in a female rat. Traces are rectified and integrated (20 ms time constant), and amplitudes are reported as percent of mean during control. Arrows indicate water infusion in the oropharynx, and vertical lines indicate swallow initiation. ANOVA showed significant differences in peak mylohyoid [*F*_(1.7, 7.9)_ = 13.1, *p* = 0.004], thyroarytenoid [*F*_(1.4, 6.5)_ = 28.2, *p* = 0.0009], and thyropharyngeus [*F*_(1.9, 8.6)_ = 13.6, *p* = 0.002], amplitudes during swallows elicited by oral water infusion. Post-hoc comparisons using the Tukey HSD test indicated that mylohyoid (33% ± 14), thyroarytenoid (68% ± 4) and thyropharyngeus (50% ± 8) amplitudes were significantly reduced during water swallows following 0.01 mg/kg buprenorphine compared to water swallows during control [mylohyoid: (95% ± 20), thyroarytenoid: (100% ± 3), thyropharyngeus: (101% ± 19), respectively. Most swallows elicited by water occurred during expiration, with no significant change in swallow breathing coordination following 0.01 mg/kg buprenorphine. D) Buprenorphine reduces frequency of swallow initiation in female rats. Frequency plot shows number of swallows elicited by oral water infusion before and after buprenorphine administration in female rats. Each circle represents the average number of swallows per stimulus for an individual animal across the buprenorphine dose response curve. ANOVA showed a significant difference in frequency of swallow occurrence [*F*_(1.8, 7.3)_ = 10.7, *p* = 0.008]. Post-hoc comparisons using the Tukey HSD test showed that significantly fewer swallows occurred following 0.01 mg/kg and 0.03 mg/kg buprenorphine. A trending decrease in swallow frequency after 0.1 mg/kg was not statistically significant (*p* = 0.09). * indicates *p* > 0.05.

Recording electrodes were placed surgically as follows: The digastric muscles were blunt dissected away from the surface of the mylohyoid, and electrodes were placed in the medial portion of the mylohyoid. Thyroarytenoid muscle electrodes were placed through the cricothyroid window into the anterior third of the vocal folds, and examined post-mortem to ensure placement accuracy. The thyropharyngeus is a fan shaped muscle that originates at the oblique line of the thyroid cartilage and courses posteriorly to the pharyngeal raphe where it meets the insertion of the contralateral thyropharyngeus. Electrodes were placed into the ventral portion of thyropharyngeus at the level of the rostral thyroid cartilage. To place electrodes in the posterior cricoarytenoid muscle, the esophagus was blunt dissected from the trachea and the trachea was elevated, which enabled direct visualization of the dorsal larynx during electrode insertion. For costal diaphragm electrode placement, the xyphoid process was palpated and elevated. Needles (1” for males, 5/8” for females) were inserted directly caudal to the sternum, and electrodes were hooked under the xyphoid process, near the costal diaphragm muscle attachment. Electrodes were placed in the left pectoralis muscle and right caudal gastrocnemius to record electrocardiogram (ECG) activity, which was used to measure heart rate and remove heart artifact from EMG traces. Correct placement of all electrodes was confirmed by visual inspection (after insertion, and post-mortem), and activation patterns during swallow and breathing, as previously published [61-65]

### Experimental Protocols

The first two experimental protocols were performed using two cohorts of male and female Sprague Dawley rats. A third protocol was performed using a cohort of female Sprague Dawley rats. A fourth protocol was performed using a cohort of male Sprague Dawley rats. 1) Cumulative buprenorphine dose response experiments were performed in 17 rats [10 male (0.5 ± 0.1 kg) and 7 female (0.24 kg ± 0.1 kg)]. 2) The 5-HT_1A_ agonist 8-OH-DPAT was administered following buprenorphine administration in 15 rats [6 male (0.7 kg ± 0.1 kg) and 9 female (0.26 kg ± 0.2 kg). 3) The partial 5-HT_1A_ agonist buspirone was given orally one hour before buprenorphine administration in 7 female rats (0.28 kg ± 0.1 kg). 4) In time control experiments for protocol 1, saline vehicle infusions were administered IV in 5 male rats (0.6 kg ± 0.1 kg). Following completion of the experimental protocol, euthanasia was induced by an overdose of sodium pentobarbital and 1 cc of saturated potassium chloride IV. In accordance withInstitutional Animal Care and Use committee guidelines, a secondary method of euthanasia was induced via pneumothorax.

#### Experiment 1: Aerodigestive Stimuli Before and After Buprenorphine Administration

Pharyngeal mechanoreceptor activation was produced by infusing 1 cc of water into the oropharynx via a 0.5-inch-long thin polyethylene catheter (outer diameter 0.5-1.0 mm) placed at the base of tongue. Esophageal mechanoreceptor activation was produced by rapidly inflating an esophageal balloon with 0.5 cc of air in less than 1 second, then maintaining inflation for 5 seconds. The balloon was attached to thin polyethylene tubing (outer diameter 0.5-1.0 mm) attached to a syringe, and placed in the upper esophagus as follows: The esophagus was blunt dissected from the trachea, and the balloon was inserted through a small incision in the caudal aspect of the thoracic esophagus and advanced proximally until just below the pharyngoesophageal segment, using the cricoid cartilage as a landmark. The esophageal catheter was secured to sternal tissue using 4-0 braided suture to ensure stable balloon placement throughout the experiment.

The following stimulation protocol was performed with at least 1 minute between each trial: 1) Water only, 2) Esophageal distension only, and 3) Esophageal distension plus water (combined stimulus), during which the esophagus was distended by balloon inflation for 5 seconds, and water was infused at the 2.5 second mark. Figure 1 displays representative swallows elicited by each stimulus.

One minute of eupnea was recorded prior to control stimulus trials. Animals then received a series of cumulative buprenorphine doses (0.01, 0.03, 0.1, and 0.3 mg/kg). Stimulus trials were repeated fifteen minutes after each dose of buprenorphine, in line with the work of Nielsen and Taylor, which demonstrates that fifteen to thirty minutes is the time to peak effect of IV buprenorphine in male Sprague Dawley rats [66]. We also recorded heart rate, respiratory rate and breathing motor patterns before and after each dose of buprenorphine.

#### Experiment 2: Systemic 8-OH-DPAT After Buprenorphine Administration

Female animals: Swallow was elicited by oral water infusion as described earlier. One minute of eupnea was recorded prior to control swallow trials. An initial dose of buprenorphine was administered (0.003 mg/kg IV), and following swallow trials, additional doses of buprenorphine were administered every fifteen minutes until opioid-induced apnea occurred (0.03 mg/kg IV among eight of nine animals in this cohort). Upon respiratory arrest, the 5-HT_1A_ agonist 8-OH-DPAT was administered (0.3 mg/kg IV). Following 8-OH-DPAT, presence or absence of respiratory effort was evaluated using EMG activity, and confirmed by direct visualization of the animal. Swallow trials were repeated eight minutes after 8-OH-DPAT in animals where breathing was restored. Finally, the competitive 5-HT_1A_ agonist was administered (1 mg/kg IV), and respiratory effort was evaluated as above. We also recorded heart rate, respiratory rate and breathing motor pattern before and after administration of each drug. 8-OH-DPAT dosage was determined by pilot experiments not included in this manuscript.

Male animals: Swallow was elicited by oral water infusion. One minute of eupnea was recorded prior to control swallow trials. Two doses of buprenorphine were administered (cumulative 0.003 and 0.03 mg/kg IV), and swallow trials were repeated fifteen minutes after each dose. Animals were subsequently treated with 8-OH-DPAT (0.3 mg/kg) and the 5HT_1A_ receptor competitive antagonist WAY-100635 (MilliporeSigma,1 mg/kg). Swallow trials were repeated eight minutes after administration of each drug. We also recorded heart rate, respiratory rate, and breathing motor pattern before and after administration of each drug.

#### Experiment 3: Buspirone Before Buprenorphine Administration

Following anesthesia with sodium pentobarbital and tracheostomy, thin polyethylene tubing (outer diameter 0.5-1.0 mm) was attached to a syringe, placed in the mouth, and advanced into the stomach. Buspirone (2.5 mg, crushed in distilled water) was then administered via orogastric gavage. One hour later, two doses of buprenorphine (cumulative 0.003 mg/kg and 0.03 mg/kg IV) were administered fifteen minutes apart. Fifteen minutes after the last dose of buprenorphine, WAY-100635 (1 mg/kg IV) was administered. We evaluated heart rate, respiratory rate, and breathing motor pattern before and after administration of each drug.

#### Experiment 4: Time Control

The following stimulation protocol was performed: 1) water only, 2) esophageal distension only, and 3) esophageal distension plus water (combined stimulus) stimulus trials as described in Experiment 1. Following control stimulus trials, four saline vehicle infusions were administered (0.5 cc IV). Stimulus trials were repeated fifteen minutes after each sham infusion. We also recorded heart rate and respiratory rate before and after each saline infusion.

### Signal and statistical analysis

EMG signals were amplified (Grass P511 AC Amplifiers, Natus Neurology), band-pass filtered (200-5000 Hz), recorded at a 10 kHz sampling rate (1401 Power3 + ADC16 Expansion, Cambridge Electronic Design), and analyzed using Spike 2 (v8, Cambridge Electronic Design). EMGs were rectified, integrated (20-ms time constant) and exported to PowerPoint (v17, Microsoft) for figure creation. Peak EMG amplitude was measured for each muscle during each swallow or breath to determine swallow and respiratory drive. For comparison across animals, raw values were normalized as the percent change in amplitude relative to the mean peak EMG amplitude during control.

Swallow-breathing coordination was evaluated by determining the phase of breathing at time of swallow initiation. Inspiration (I) was defined as the period from the onset of breathing-related diaphragm activity to the peak of the diaphragm burst. Expiration (E) was defined as the period from peak diaphragm activity to the onset of subsequent diaphragm activation, and further subdivided into early expiration (E1; the period from peak diaphragm amplitude to diaphragm quiescence) and late expiration (E2; the period from offset of diaphragm activity to the onset of subsequent diaphragm activity) [14, 54]. Swallows that occurred during inspiration, early expiration and late expiration were coded as 1, 2 and 3 respectively, and a Wilcoxon signed-rank test was used to evaluate differences in swallow-breathing coordination across conditions.

The primary outcome measures are amplitudes of submental, laryngeal, pharyngeal, and inspiratory electromyograms (EMGs), and frequency of swallow occurrence. Secondary outcome measures included respiratory and heart rate. GraphPad Prism statistical software (GraphPad Software, Boston, MA) was used to perform analysis of variance (ANOVA) and Tukey HSD post-hoc comparisons. Data expressed as mean ± standard deviation. For all measures, a difference was considered significant if the *p*-value was less than 0.05.

## Results

First, we evaluated the effects of varied upper aerodigestive stimulation on oropharyngeal swallow initiation and motor pattern. Figure 1 demonstrates that oral water infusion, rapid esophageal distension, and esophageal distension plus water (combined stimulus) reliably elicit swallow, and that swallow-related mylohyoid (laryngeal elevator) and thyroarytenoid peak amplitudes are larger during swallows elicited by esophageal distension plus water. There was no difference in swallow-related thyropharyngeus activity across stimulus conditions.

Second, we assessed the effects of systemic buprenorphine administration on oropharyngeal swallow in male and female rats. Figure 2 demonstrates that buprenorphine reduces swallow-related laryngeal elevation in male rats (Fig. 2A), and reduces swallow-related laryngeal elevation, laryngeal adduction and pharyngeal constriction in female rats. In contrast to female animals, thyroarytenoid (laryngeal adductor) and thyropharyngeus (pharyngeal constrictor) activity during swallow were not significantly affected by buprenorphine in males. Frequency of swallow initiation was reduced in both male and female animals following buprenorphine administration (Fig. 2B, 2D). To determine if these changes in function were due to effects of buprenorphine or effects of time, we performed a series of time control experiments (*n* = 5) in which animals received IV saline vehicle infusions instead of buprenorphine. There were no significant changes in swallow-related peak amplitude (mylohyoid, thyroarytenoid, thyropharyngeus), or swallow frequency (Fig. 2B, 2D) across stimulus conditions (water, esophageal distension, esophageal distension plus water) and saline infusions among the time control group.

Third, we evaluated the effects of systemic buprenorphine administration on breathing and survival in male and female rats. Figure 3 shows that while buprenorphine had no significant effect on respiratory rate in male animals (Fig. 3B), respiratory rate was reduced following buprenorphine administration in female animals (Fig. 3E). Female animals had a lower threshold for opioid-induced apnea (Fig. 3F), and there was a difference in the survival distributions of male and female animals along the buprenorphine dose response curve (Fig. 3G).

**Figure 3.**
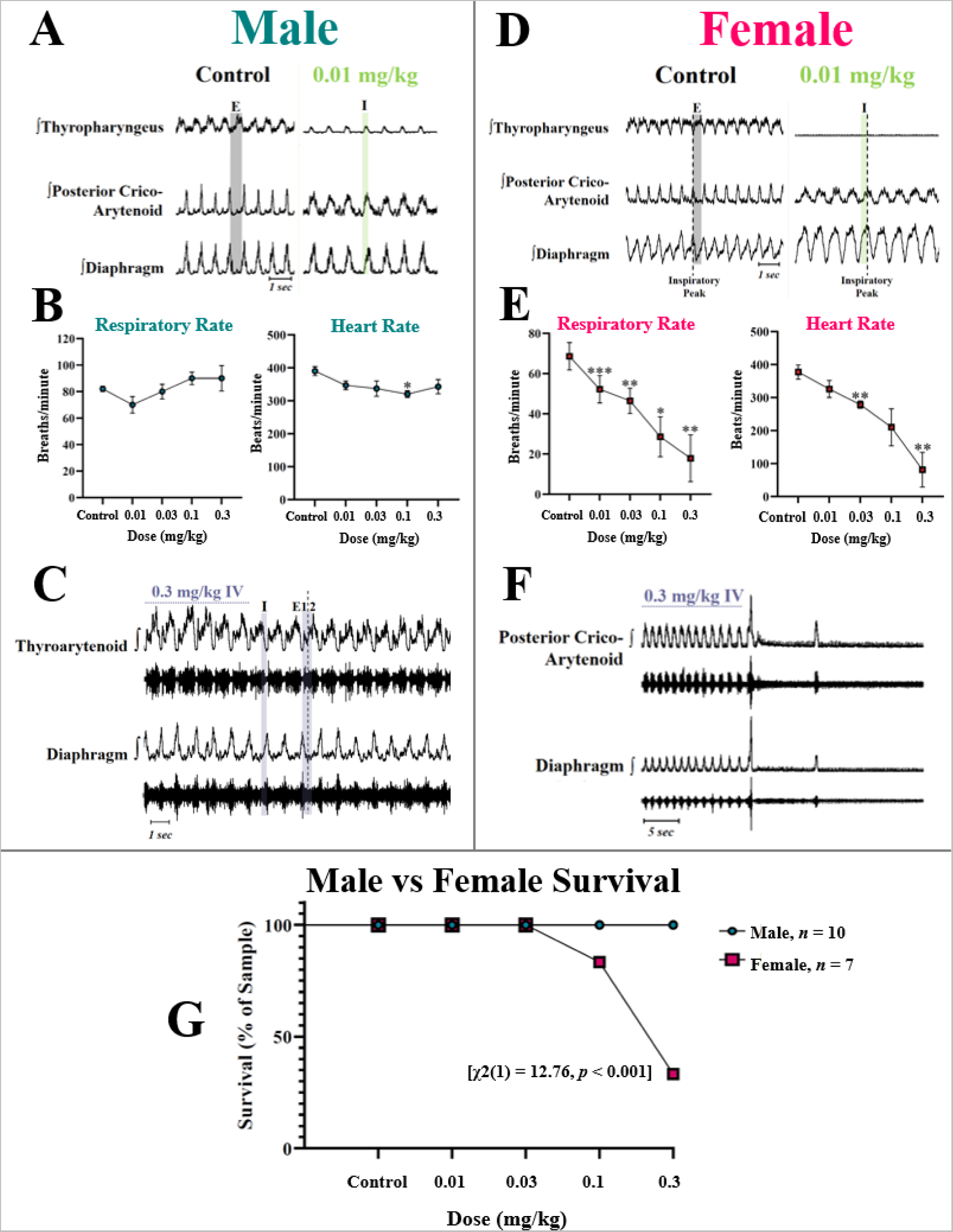
The opioid buprenorphine disproportionately depresses respiration in female rats. A) Representative electromyogram (EMG) traces show thyropharyngeus, posterior cricoarytenoid, and diaphragm muscle activity during eupnea before and after buprenorphine administration (0.01 mg/kg IV) in a male rat. Traces are rectified and integrated (20 ms time constant). The thyropharyngeus (inferior pharyngeal constrictor) is active during both breathing and swallow. During control, the thyropharyngeus demonstrates phasic expiratory activity during breathing (gray bar). After buprenorphine, the activity of the thyropharyngeus shifts and is phasic during inspiration (green bar). A Wilcoxon signed-rank test detected a significant change in thyropharyngeus respiratory phase preference, with significantly more male animals demonstrating inspiratory activity following buprenorphine administration (*Z* = -2.7, *p* = 0.004).B) Plots show mean respiratory rate and heart rate among male rats across a cumulative buprenorphine dose response curve (0.01, 0.03, 0.1 and 0.3 mg/kg IV). Plotted values represent mean breaths per minute and beats per minute, respectively. Buprenorphine did not significantly impact respiratory rate in male animals. Analysis of variance (ANOVA) showed a significant difference in heart rate [*F*_(2.5, 19.2)_ = 3.8, *p* = 0.03], and post-hoc comparisons using the Tukey HSD test indicated heart rate was significantly reduced following 0.1 mg/kg buprenorphine (320 ± 30) compared to control (390 ± 42.4). * indicates *p* < 0.05. C) Representative EMG traces showing thyroarytenoid and diaphragm activity during eupnea following high dose buprenorphine (0.3 mg/kg IV) in a male rat. Traces are rectified and integrated (20 ms time constant). Horizontal dotted line indicates IV infusion. Vertical dotted line demarcates the beginning of diaphragm quiescence within a single breath cycle, which indicates the end of early expiration (E1) and the beginning of late expiration (E2). The thyroarytenoid (laryngeal adductor) is active during both breathing and swallow, and normally demonstrates phasic expiratory activity during eupnea. Traces depict stable respiratory activity in the upper airway and diaphragm after high dose buprenorphine in a male animal. D) Representative electromyogram (EMG) traces show thyropharyngeus, posterior cricoarytenoid, and diaphragm activity during eupnea before and after buprenorphine administration (0.01 mg/kg IV) in a female rat. Traces are rectified and integrated (20 ms time constant). Vertical dotted lines indicate peak diaphragm amplitude of a single breath, which marks the end of inspiration (I) and the beginning of early expiration (EI). During control, the thyropharyngeus is active during expiration (highlighted in gray). Breathing-related thyropharyngeus activity is lost following buprenorphine administration. Five female animals demonstrated reduced breathing-related thyropharyngeus activity after buprenorphine (0.01 mg/kg IV), but the effect was not significant as a group. Representative trace depicts increased breathing-related diaphragm activation following buprenorphine administration. Diaphragm EMG amplitude increased in 4 of 7 female animals (Control: 100% ± 0; 0.01mg/kg: 103% ± 22) and 9 of 10 male animals (Control: 100% ± 0; 0.01 mg/kg: 119% ± 51) after buprenorphine administration. Diaphragm burst duration increased in all female animals (Control: 339 ms ± 122; 0.01 mg/kg: 425 ms ± 151) and 9 of 10 male animals (Control: 258 ms ± 56; 0.01 mg/kg: 340 ms ± 66) after buprenorphine administration. ANOVA detected no statistical differences in diaphragm amplitude or duration after buprenorphine in the male or female groups, however the observed increase in average values may be physiologically meaningful. We speculate that in the setting of opioid-induced respiratory failure, reduced laryngeal resistance during expiration may contribute to increased amplitude and duration of diaphragm EMG amplitude during inspiration. E) Plots show mean respiratory rate and heart rate among female rats across a cumulative buprenorphine dose response curve (0.01, 0.03, 0.1, and 0.3 mg/kg IV). Plotted values represent mean breaths per minute and beats per minute, respectively. ANOVA showed a significant difference in respiratory rate among female animals [*F*_(1.5, 6.3)_ = 11, *p* = 0.01], and post-hoc comparisons using the Tukey HSD test showed a significant decrease in respiratory rate following 0.01 mg/kg (52 ± 18), 0.03 mg/kg (46 ± 17), 0.1 mg/kg (29 ± 26), and 0.3 mg/kg buprenorphine (18 ± 31) compared to control (69 ± 18). ANOVA showed a significant difference in heart rate among female animals, and post-hoc comparisons using the Tukey HSD test showed a significant decrease in heart rate following 0.03 mg/kg (279 ± 29) and 0.3 mg/kg buprenorphine (81 ± 139) compared to control (377 ± 57). *** indicates *p* < 0.001, ** indicates *p* < 0.01, and * indicates *p* < 0.05. F) Representative EMG traces showing posterior cricoarytenoid and diaphragm activity during breathing, and apnea following high dose buprenorphine (0.3 mg/kg IV) administration in a female rat. Traces are rectified and integrated (20 ms time constant). The posterior cricoarytenoid (laryngeal abductor) demonstrates phasic activity during eupnea, and functions to open the glottis during inspiration. Traces depict loss of inspiratory effort in the upper airway and diaphragm following high dose buprenorphine. Breathing did not recover in this female animal. G) Plot of survival proportions among male and female rats following buprenorphine administration (cumulative 0.01, 0.03, 0.1 and 0.3 mg/kg IV). A log rank test detected a significant difference in the survival distribution between males and females. More than 70% of female animals succumbed to respiratory arrest following high dose buprenorphine (0.1 and 0.3 mg/kg), while all male animals maintained stable respiratory effort across doses.

Fourth, we tested the utility of the 5-HT_1A_ agonist 8-OH-DPAT in the restoration of breathing following buprenorphine-induced apnea in female animals. Systemic administration of 8-OH-DPAT (0.3 mg/kg IV) restored breathing following respiratory arrest in 75% of female animals, and this effect was reversed by the competitive 5-HT_1A_ antagonist WAY-100635 (Fig. 4A). Unlike female animals, male animals maintained regular respiratory effort across all conditions (Fig. 5B, 5C).

**Figure 4.**
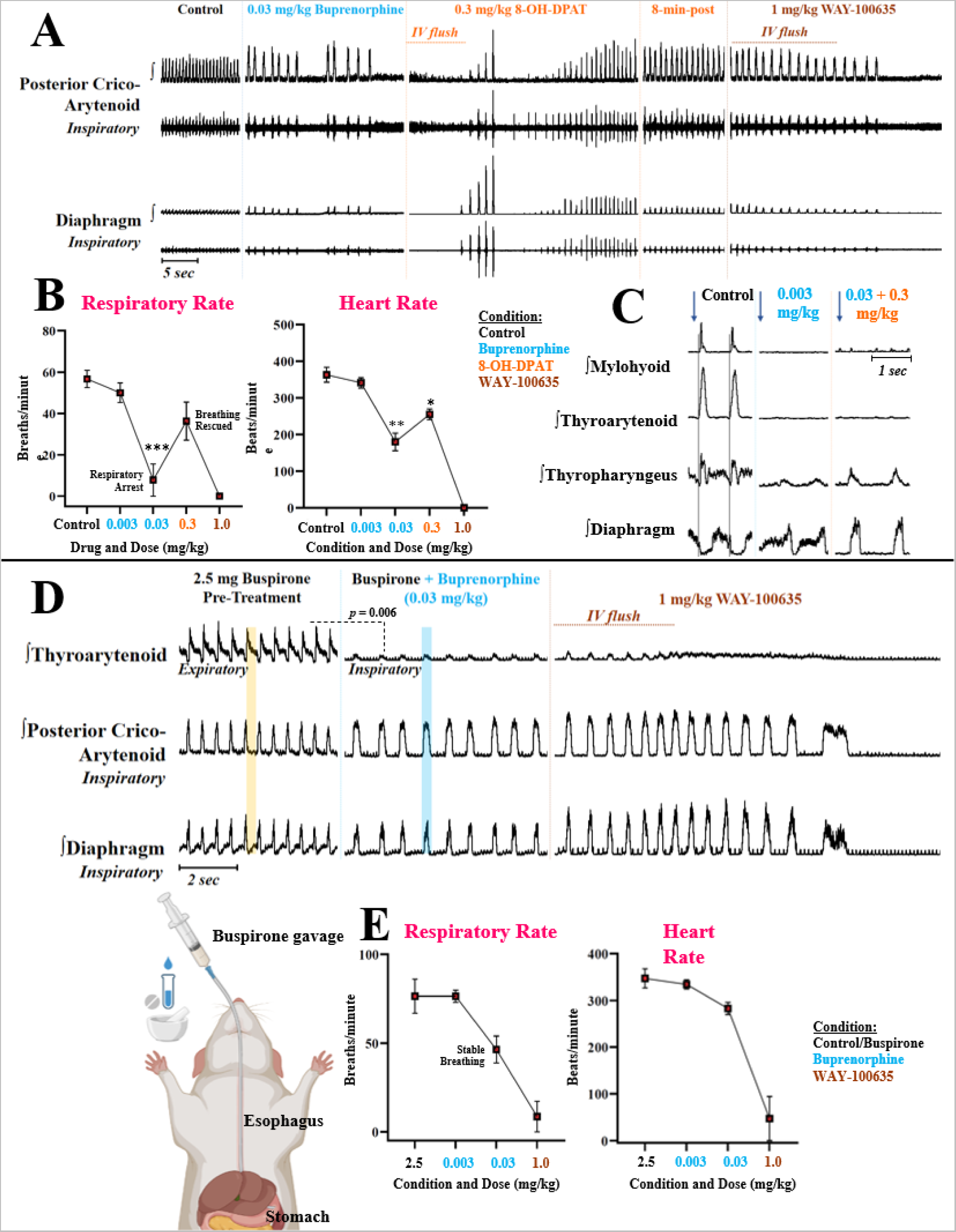
5-HT1_A_ agonists restore and preserve breathing following buprenorphine administration in female rats. A) To test the hypothesis that systemic administration of a 5-HT_1A_ agonist would restore breathing following opioid induced apnea, we performed experiments in *n* = 9 freely breathing pentobarbital anesthetized adult female Sprague Dawley rats. A) Representative traces from posterior cricoarytenoid (laryngeal abductor) and diaphragm electromyograms (EMGs) show activity during breathing in a female rat. Traces are rectified and integrated (20-ms). Following a period of eupnea (control), buprenorphine was titrated IV until apnea occurred (0.03 mg/kg in this animal). The 5-HT_1A_ agonist 8-OH-DPAT was then administered (0.3 mg/kg IV), and inspiratory effort was restored within 30 seconds of infusion. Breathing remained stable for over eight minutes following 8-OH-DPAT, and was abolished by administration of the competitive 5-HT_1A_ antagonist WAY-100635 (1 mg/kg IV). Eight of nine animals stopped breathing following buprenorphine administration (one animal resisted opioid induced apnea). 8-OH-DPAT restored breathing in six animals (no restoration in two animals), and this effect was reversed by WAY-100635. B) Plots show mean respiratory rate and heart rate among female rats after IV administration of buprenorphine, 8-OH-DPAT, and WAY-100635. Plotted values represent mean breaths per minute and beats per minute, respectively. Analysis of variance (ANOVA) detected significant differences in respiratory rate [*F*_(2, 13.8)_ = 18.7, *p* = 0.0001] and heart rate [*F*_(2.6, 21.2)_ = 41.6, *p* < 0.0001]. Post-hoc comparisons using the Tukey HSD test indicated that respiratory rate was significantly reduced following 0.03 mg/kg buprenorphine (8 ± 30) compared to control (57 ± 13). Respiratory rate increased in seven of nine animals following 0.3 mg/kg 8-OH-DPAT (compared to 0.03 mg/kg buprenorphine), but this effect was not statistically significant as a group (*p* = 0.09). Post-hoc comparisons using the Tukey HSD test indicated that heart rate was significantly reduced following 0.03 mg/kg buprenorphine (180 ± 72) compared to control (363 ± 56), and significantly increased following 0.3 mg/kg 8-OH-DPAT (255 ± 29) compared to 0.03 mg/kg buprenorphine. *** indicates *p* < 0.001, ** indicates *p* < 0.01, *p* < 0.05. C) Representative traces from mylohyoid (laryngeal elevator), thyroarytenoid (laryngeal adductor), thyropharyngeus (pharyngeal constrictor) and diaphragm EMGs show response to oral water infusion before and after administration of buprenorphine (0.03 mg/kg IV) and 8-OH-DPAT (0.3 mg/kg IV) in a female rat. Traces are rectified and integrated (20 ms time constant). Arrows indicate oral water infusion and vertical lines indicate swallow onset. Swallow initiation was suppressed following buprenorphine administration (0.03 mg/kg IV). 8-OH-DPAT administration did not restore responsiveness to oral water infusion following total swallow suppression by buprenorphine in female animals. D) To test the hypothesis that pre-treatment with the partial 5-HT_1A_ agonist buspirone would prevent opioid induced apnea in females, experiments were performed in *n* = 7 adult female Sprague Dawley rats. Representative traces from thyroarytenoid (laryngeal adductor), posterior cricoarytenoid (laryngeal abductor) and diaphragm EMGs show activity during eupnea in a female rat pre-treated with buspirone (2.5 mg orogastric gavage) and subsequently treated with buprenorphine (0.03 mg/kg IV) and WAY-100635 (1 mg/kg IV). Traces are rectified and integrated (20 ms time constant), and amplitudes are reported as percent mean of control. Yellow rectangle highlights the expiratory phase of one respiratory cycle. Blue rectangle highlights the inspiratory phase of one respiratory cycle. Representative trace of thyroarytenoid activity depicts phasic expiratory activity during buspirone control, and reduced peak amplitude, phasic inspiratory activity, and expiratory quiescence following buprenorphine administration. ANOVA detected significant differences in breathing-related thyroarytenoid amplitude across conditions [*F*_(1.3, 7.6)_ = 26.3, *p* = 0.0007], and post-hoc comparisons using the Tukey HSD test indicated a significant decrease in breathing-related thyroarytenoid amplitude following 0.03 mg/kg buprenorphine (66% ± 18) compared to control (100% ± 0). A Wilcoxon signed-rank test detected a significant difference in laryngeal respiratory phase preference, with significantly more animals demonstrating phasic inspiratory thyroarytenoid activity following buprenorphine administration compared to control, when phasic thyroarytenoid activity is expiratory (*z* = -1.86, *p* = 0.03). EMG traces depict stable respiratory effort after a dose of buprenorphine that produced apnea in animals that did not receive buspirone (0.03 mg/kg IV, see panel A), and cessation of breathing following administration of the competitive 5-HT_1A_ antagonist WAY-100635. All seven animals pre-treated with buspirone maintained regular breathing across buprenorphine doses, and breathing was abolished in six animals following WAY-100635 (one animal resisted respiratory arrest). E) Plots show respiratory rate and heart rate among female rats pre-treated with buspirone following administration of buprenorphine (cumulative 0.003 and 0.03 mg/kg IV) and WAY-100635. Plotted values represent mean breaths per minute and beats per minute, respectively. There was no significant change in respiratory rate or heart rate following buprenorphine administration among female animals pre-treated with buspirone, compared to control. A log rank test detected a significant difference in survival between experimental groups, with higher median survival following buprenorphine administration among female animals pre-treated with buspirone than animals that did not receive buspirone [χ2(1) = 11.67, *p* < 0.001].

**Figure 5.**
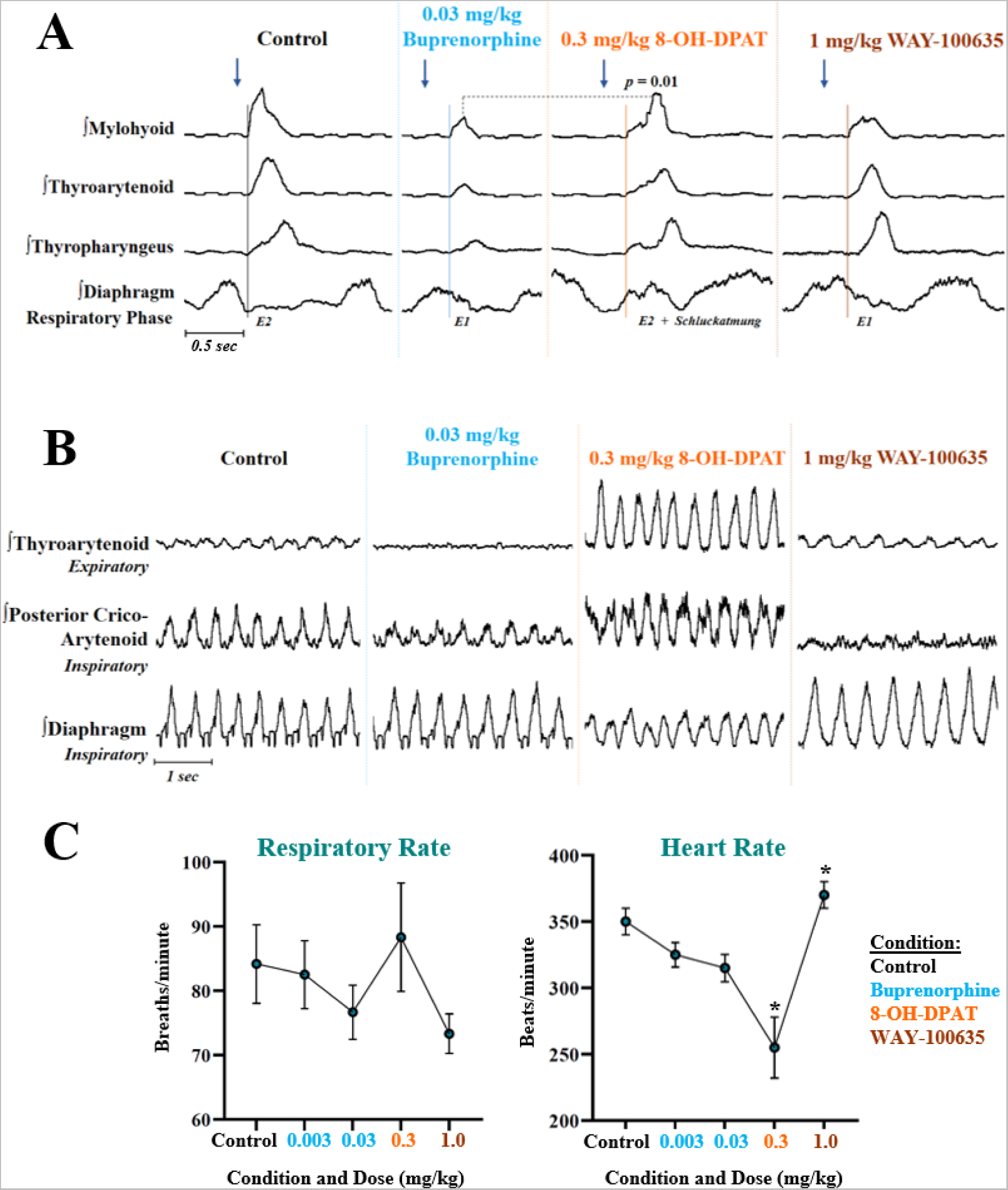
The 5-HT1_A_ agonist 8-OH-DPAT restores laryngeal elevation during swallow following buprenorphine administration in male rats. To test the hypothesis that systemic 8-OH-DPAT administration would result in correction toward baseline swallow measures following buprenorphine-induced alteration of swallow motor pattern, experiments were performed in *n* = 6 spontaneously breathing pentobarbital anesthetized adult male Sprague Dawley rats. A) Representative electromyogram (EMG) traces show mylohyoid (laryngeal elevator), thyroarytenoid (laryngeal adductor), thyropharyngeus (pharyngeal constrictor) and diaphragm muscle activity during swallow following buprenorphine (0.03 mg/kg IV), 8-OH-DPAT (0.3 mg/kg IV) and WAY-100635 (1 mg/kg IV) administration. Traces are rectified and integrated (20 ms time constant), and amplitudes are reported as percent mean of control. Arrows indicate oral water infusion and vertical lines indicate swallow onset. Analysis of variance (ANOVA) showed a significant difference in peak mylohyoid amplitude during swallow across conditions [*F*_(1.7, 6.9)_ = 19.3), *p* = 0.002], and post-hoc comparisons using the Tukey HSD test revealed a significant decrease in peak mylohyoid amplitude following 0.03 mg/kg buprenorphine (51% ± 18) compared to control (100% ± 0), and a significant increase in peak mylohyoid amplitude following 0.3 gm/kg 8-OH-DPAT (88% ± 22) compared to 0.03 mg/kg buprenorphine. Following administration of the competitive 5-HT_1A_ antagonist WAY-100635, swallow-related mylohyoid amplitude decreased in five of six animals (63% ± 34), and was no longer significantly different than mylohyoid amplitude following 0.03 mg/kg buprenorphine (*p* = 0.8). A Wilcoxon signed-rank test detected a significant change in swallow breathing coordination, with significantly more swallows occurring during early expiration (E1) following 0.03 mg/kg buprenorphine (*z* = -3.3, *p* = 0.004) compared to control, when most swallows occurred during late expiration (E2). This effect persisted following 0.3 mg/kg 8-OH-DPAT (*z* = -2.7, *p* = 0.0004) and WAY-100635 (*z* = -3.1, *p* = 0.001), with significantly more swallows initiated during early expiration (E1) at the group level. ANOVA detected no difference in time-to-peak of swallow-related mylohyoid activation [*F*_(2, 8 = 1.46)_, p = 0.27)], indicating that the order of muscle fiber recruitment was unchanged following buprenorphine and 8-OH-DPAT administration. B) Representative electromyogram (EMG) traces of breathing-related thyroarytenoid (laryngeal adductor), posterior cricoarytenoid (laryngeal abductor) and diaphragm activity during control and following IV administration of buprenorphine (0.03 mg/kg), 8-OH-DPAT (0.3 mg/kg), and WAY-100635 (1 mg/kg) in a male rat. Traces are rectified and integrated (50 ms time constant). Traces depict reduction in breathing-related activation of laryngeal muscles (thyroarytenoid and posterior cricoarytenoid) following buprenorphine (0.03), and an increase in laryngeal drive during breathing following 8-OH-DPAT (0.3 mg/kg) that is reversed by the competitive 5-HT_1A_ agonist WAY-100635. This series of effects occurred in four of six animals, but was non-significant as a group. C) Plots show respiratory rate and heart rate among male rats following IV administration of buprenorphine (0.003 and 0.03 mg/kg), 8-OH-DPAT (0.3 mg/kg), and WAY-100635 (1 mg/kg). Plotted values represent mean breaths per minute and beats per minute, respectively. There were no significant differences in respiratory rate among male rats across doses. ANOVA indicated significant differences in heart rate across conditions, and post-hoc comparisons using the Tukey HSD test showed a significant decrease in heart rate following 8-OH-DPAT (255 ± 62) compared to control (350 ± 27), and a significant increase in heart rate following WAY-100635 (370 ± 27) compared to 8-OH-DPAT. Unlike female animals, all male animals maintained stable respiratory effort across conditions. * indicates *p* < 0.05.

Fifth, we wanted to determine if pre-treatment with oral buspirone (a partial 5-HT_1A_ agonist) would prevent buprenorphine-induced apnea in female animals. Animals that received 2.5 mg oral buspirone maintained stable respiratory effort following doses of buprenorphine that produced apnea in animals that did not receive buspirone, and demonstrated significantly higher median survival following opioid administration (Fig. 4D, 4E).

Finally, we used systemic 8-OH-DPAT to evaluate the utility of 5-HT_1A_ agonists in the restoration of swallow function following opioid-induced swallow depression. Swallow initiation was not recovered by IV 8-OH-DPAT following total suppression of swallow post-buprenorphine in female animals (Fig. 4C). However, among male animals that still initiated swallow, mylohyoid (laryngeal elevator) amplitude was reduced after buprenorphine administration, with correction toward baseline following IV 8-OH-DPAT (Fig. 5A). There were no changes in swallow-related thyroarytenoid or thyropharyngeus amplitude across conditions in male animals.

## Discussion

Swallow and breathing are vulnerable to depression by opioids, even following low doses of the partial opioid agonist buprenorphine. Profound differences in sensitivity to buprenorphine between male and female rats indicate need for further evaluation of sex differences in humans. Our finding that 5-HT_1A_ agonists improve swallow and breathing measures following opioid administration expands on previous work in rats [52, 67]. We advocate that these results warrant further translational studies to enable successful application to human medicine.

### Buprenorphine Disproportionately Depresses Swallow and Breathing in Female Rats

A functional aerodigestive tract is dependent upon central integration of feedback from several afferent beds in the larynx, pharynx and esophagus [7, 8, 68, 69]. Our results support our first hypothesis, that swallow motor pattern is modulated based on the location and intensity of peripheral aerodigestive stimulation. Following buprenorphine administration, pharyngeal swallow function was depressed, and laryngeal accommodation to a maximal stimulus was lost. Buprenorphine blunted aerodigestive responses such that often, no swallow or protective response was elicited by stimuli that effectively flooded the upper airway. When swallow did occur post-buprenorphine, there was significant reduction of EMG amplitudes, and altered swallow-breathing coordination. While swallow was depressed in both male and female animals, swallow was frankly depressed by buprenorphine in females. These findings support our second hypothesis, that the opioid buprenorphine would depress swallow function.

The swallowing musculature is driven by anatomically distinct motor nuclei: The mylohyoid is innervated by the trigeminal nerve (CN V); the thyroarytenoid is innervated by the recurrent laryngeal branch of the vagus nerve (CN X); and the thyropharyngeus is innervated by the glossopharyngeal and vagus nerves (CN IX, X) [57, 58, 68]. We hypothesize that buprenorphine centrally depresses swallow through its actions on opioid receptors throughout the brainstem swallow pattern generator. Buprenorphine is a partial mu-opioid agonist, as well as a kappa-opioid inverse agonist and delta-opioid antagonist [70]. Mu-opioid receptors have been identified in several brainstem regions, including the nucleus tractus solitarius (NTS), which is the site of initiation of the central swallow command [16, 17, 20, 23]. Relatively few studies have investigated kappa-and delta-opioid receptors in the rat brainstem, and potential actions of these receptors on regions important for swallow and breathing are unclear [71, 72]. Our findings indicate that buprenorphine disrupts neural networks throughout the brainstem resulting in dysphagia, but more work is needed to determine the molecular basis and site of action of buprenorphine’s effects on the swallow pattern generator.

The activity of each swallow-related muscle we measured represents a critical component of pharyngeal swallow: The mylohyoid (laryngeal elevator) helps lift the larynx above food or liquid as it passes through the pharynx; the thyroarytenoid (laryngeal adductor) functions to seal the larynx from ingested material; and the thyropharyngeus (pharyngeal constrictor) aids bolus clearance into the esophagus [6, 10, 61]. Clinically, even slight disturbances of aerodigestive regulation can result in aspiration-related complications (e.g., aspiration pneumonia, death) [1, 12, 63, 73-75]. Alterations in swallow-related EMG amplitude following buprenorphine administration were predictive of severe dysphagia, most remarkably in female animals.

Previous studies have reported that buprenorphine-induced respiratory depression has a ceiling effect [41, 43]. Our third hypothesis, that buprenorphine would not produce dose-dependent respiratory depression, was refuted by experiments performed in female rats. These experiments demonstrated progressive respiratory depression and respiratory arrest as buprenorphine dosage increased. We obtained a different result in male rats, which showed no significant slow in respiratory rate following large doses of buprenorphine. Our finding that buprenorphine powerfully depresses swallow, and produces terminal respiratory arrest in female animals, suggests that even opioids considered to be safe can negatively affect survival.

To our knowledge, this study is the first to evaluate sex-specific effects of buprenorphine on breathing and swallowing. Our finding that breathing and swallowing are regulated differently in male and female animals is consistent with a limited, but growing number of studies dedicated to physiological sex differences. Studies by Huff and colleagues [76, 77] demonstrated evidence of sex-specific modulation of breathing and swallowing when a mechanical challenge or airway anesthesia was experimentally induced in rats. A recent study comparing opioid-induced respiratory depression between male and female rats found that females demonstrated a greater degree of heroin-induced respiratory depression than males [78, 79]. The few studies performed in humans have also shown that women are more sensitive to morphine-induced respiratory depression than men. These reports, and the present results, highlight the importance of sex differences in the study of control of breathing and its depression by opioids.

### 5-HT_1A_: A Promising Target for Opioid-Induced Depression of Breathing and Swallowing

Serotonin is a modulatory neurotransmitter synthesized in the brainstem raphe nuclei that has been shown to modulate breathing and swallow [44, 48, 49, 80]. The effects of serotonin are complex, and depend on the anatomical distribution of receptor subtypes on a given neuron or area [49]. The 5-HT_1A_ receptor is an inhibitory serotonin receptor subtype that is distributed throughout the brainstem and has been implicated in regions important for breathing and swallow [45, 46, 50, 51, 67, 80].

Experimentally, activation of medullary 5-HT_1A_ receptors has been shown to stimulate respiration [50, 51, 81]. Previous studies in rats have shown that 5-HT_1A_ agonists restore respiratory function following opioid-induced respiratory depression and arrest in males [52, 67, 81]. Our experiments in female rats support our fourth and fifth hypotheses, that 8-OH-DPAT would restore breathing following opioid-induced apnea, and buspirone would preserve breathing following buprenorphine administration. Of note, a clinical trial in humans was performed, and concluded that buspirone does not antagonize opioid-induced respiratory depression based on CO_2_ rebreathing following morphine administration in healthy subjects [26].

We do not claim that 5-HT_1A_ agonists restore respiratory function to normal range following opioid administration. However, our results do show that 8-OH-DPAT and buspirone extend survival by countering terminal respiratory arrest. We have replicated and expanded upon the results presented by Sahibzada and colleagues, and advocate for future research aiming to translate these findings to human medicine [52].

5HT_1A_ agonists have been shown to improve dysphagia symptoms and esophageal motility in humans [26, 82, 83]. Following total depression of breathing and swallowing in female rats, 8-OH-DPAT restored respiratory rhythmicity, but did not restore excitability of the swallow pattern generator. As we found in our initial dose-response experiments, male animals that were still stimulable for swallow following buprenorphine demonstrated significantly reduced mylohyoid amplitude. This decline in mylohyoid activation was reversed by subsequent 8-OH-DPAT administration. These findings support our hypothesis that 8-OH-DPAT would measurably improve buprenorphine-induced dysphagia, with the caveat that 8-OH-DPAT enhanced submental neural drive during swallow, but did not restore excitability of the swallow reflex initiation following total suppression.

Despite reports from several groups that 5-HT_1A_ agonists stimulate breathing, the mechanisms through which inhibitory 5-HT_1A_ receptors modulate central pattern generators have not been definitively described [50-52, 67, 81]. It has been proposed that following opioid-induced depression of respiratory frequency, activation of 5-HT_1A_ receptors stimulates respiration by disinhibiting post-inspiratory neurons [81]. It is possible that a different action is responsible for restoration of breathing following opioid-induced respiratory arrest, but more work is needed to elucidate the mechanism. We can attribute the effects we observed with buspirone and 8-OH-DPAT to action on 5-HT_1A_ receptors, as protective effects on breathing and swallowing were reversed by the competitive 5-HT_1A_ agonist WAY-100635.

### Clinical Impact

There are currently no pharmacological treatments for pharyngeal dysphagia, a diagnosis that is challenging to treat and associated with poor outcomes [74, 75, 84-89]. There is need for a drug that stabilizes breathing for patients who are prescribed opioids and have risk factors for respiratory depression. Although naloxone effectively reverses opioid overdose, it is not a suitable concomitant therapy because it also reverses antinociception, and can precipitate withdrawal [23, 78]. Our pre-clinical data show that 5-HT_1A_ is a promising receptor for drug development aimed at treating dysphagia and opioid-induced respiratory arrest. Few 5-HT_1A_ agonists are currently available for use in humans, but buspirone is an FDA-approved anxiolytic with a small side effect profile that may also be employed to treat disorders of breathing and swallowing [90]. More research is needed on opioids, serotonin and the neural networks controlling breathing and swallowing in order to develop effective pharmacological therapies for humans.

## Acknowledgments

Figures were created with BioRender.

## Disclaimers

The funders had no role in study design, data collection and analysis, decision to publish, or preparation of the manuscript.

## Author Contributions

M.F., K.I., and T.P. conceived and designed research; M.F., I.M., N.F, and T.P. performed experiments; M.F., K.I., and T.P. analyzed data; M.F., N.M., K.I., and T.P. interpreted results of experiments; M.F., K.I., and T.P. prepared figures; M.F. drafted manuscript; M.F., N.M., K.I., and T.P. edited and revised manuscript; M.F., I.M., N.F., N.M, K.I., and T.P. approved final version of manuscript.

